# GABA, Glutamate dynamics and BOLD observed during cognitive processing in psychosis patients with hallucinatory traits

**DOI:** 10.1101/2024.12.13.628297

**Authors:** Alexander R. Craven, Gerard Dwyer, Lars Ersland, Katarzyna Kazimierczak, Lin Lilleskare, Ralph Noeske, Lydia Brunvoll Sandøy, Erik Johnsen, Kenneth Hugdahl

## Abstract

The perception of a voice in the absence of an external auditory source – an auditory verbal hallucination – is a characteristic symptom of schizophrenia. To better understand this phenomenon requires integration of findings across behavioural, functional, and neurochemical levels. We address this with a locally adapted MEGA-PRESS sequence incorporating interleaved unsuppressed water acquisitions, allowing concurrent assessment of behaviour, blood-oxygenation-level-dependent (BOLD) functional changes, Glutamate+Glutamine (Glx), and GABA, synchronised with a cognitive (flanker) task. We acquired data from the anterior cingulate cortex (ACC) of 51 patients with psychosis (predominantly schizophrenia spectrum disorder) and hallucinations, matched to healthy controls. Consistent with the notion of an excitatory/inhibitory imbalance, we hypothesized differential effects for Glx and GABA between groups, and aberrant dynamics in response to task. Results showed impaired task performance, lower baseline Glx and positive association between Glx and BOLD in patients, contrasting a negative correlation in healthy controls. Task-related increases in Glx were observed in both groups, with no significant difference between groups. No significant effects were observed for GABA. These findings suggest that a putative excitatory/inhibitory imbalance affecting inhibitory control in the ACC is primarily observed as tonic, baseline glutamate differences, rather than GABAergic effects or aberrant dynamics in relation to a task.

**Highlights:**

- In-vivo, GABA-edited functional ^1^H-MRS data were collected from 51 patients with hallucinations and a similar number of matched healthy controls
- Reduced Glutamate+Glutamine (Glx) levels were observed in the patient group.
- BOLD association to baseline Glutamate+Glutamine (Glx) differed between patients and controls
- Robust task-related increases in measured Glx were observed in the Anterior Cingulate Cortex (ACC)
- Task-related changes in measured Glx did not differ between patients and controls

## 1 Introduction

The perception of a voice in the absence of an external auditory source – an auditory verbal hallucination (AVH) – is a characteristic symptom of schizophrenia, manifesting at different “levels of explanation” [1,2]. These range from broad, high-level, macro-scale aspects of cultural and social context, through to clinical symptoms and diagnoses, cognitive factors, and increasingly intricate, mechanistic aspects of integrated neural systems and networks, individual synapses and neurotransmitters forming a part of those systems, and the molecular processes occurring therein. While observations at all levels have been informative in guiding targets for treatment and for further research, these when considered in isolation have not yielded a comprehensive explanation of the complex, multi-faceted phenomenon. Indeed, without effective vertical integration to harmonise findings across levels there is a risk that the theories and models under investigation become dissociated from the very phenomena which they attempt to explain[1].

Amongst current theories aiming to explain the phenomenology of patients “hearing voices” is a putative breakdown of the dynamic interplay between bottom-up (excitatory, perceptual) and top-down (inhibitory control) cognitive processes, perhaps reflected in the dynamic interaction of excitatory and inhibitory neurotransmitters[3]: glutamate (Glu) and γ- aminobutyric acid (GABA) respectively. While there are published imaging and static spectroscopy findings to support this model[2,4], there is currently limited data to bridge clinical and imaging findings with neurotransmitter dynamics at the receptor level, due in part to the challenges associated with reliably measuring these neurotransmitters in a dynamic (functional) setting.

Acquisition and reliable analysis of functional Magnetic Resonance Spectroscopy (fMRS) data is challenging for two reasons: firstly, the metabolite signals of interest are several orders of magnitude weaker than the water signal which forms the basis of blood-oxygenation level dependent (BOLD) functional Magnetic Resonance Imaging (fMRI) measurement, leading to a substantial trade-off against spatial and temporal resolution.

Furthermore, BOLD-related changes in signal relaxation and local shim quality have a direct impact on spectral line-shape which may affect the quantification of certain metabolites.

While this represents a potential confound for metabolite quantification[5], it also provides an opportunity for simultaneously assessing task-induced BOLD dynamics from unsuppressed water data obtained during the MRS acquisition[6,7]. Extending the approach of Apšvalka et al [8] to a GABA-editing (MEGA-PRESS) context[9,10], we previously demonstrated a technique for simultaneously obtaining time-resolved, GABA-edited, MRS data and an indication of local BOLD response[11], incorporating unsuppressed water reference signals at a regular interval within the edited acquisition scheme. The GABA-editing technique yields estimates for GABA with some contribution from underlying co-edited signals (GABA+), and estimates for Glutamate and Glutamine combined (Glx). The present study applies the same acquisition and analysis techniques (and a matched subset of the same healthy controls) in a case-control context with psychosis patients, predominantly with a diagnosed schizophrenia spectrum disorder (SSD).

Consistent with the notion of an excitatory/inhibitory imbalance, we hypothesize differential effects for Glx and GABA between healthy controls and patients, and aberrant dynamics in one or both in relation to a cognitive task. Furthermore, based on the findings of Falkenberg et al [12], we anticipate positive correlation between BOLD-fMRI activation and baseline Glx in the patient group, with decreased Glx levels associated with impaired executive control functioning in that group. We further anticipate a negative correlation between BOLD-fMRI activation and baseline Glx in the healthy controls.

## 2 Methods

### 2.1 In-vivo data collection

#### 2.1.1 Subject recruitment and demographics

The study included 54 psychiatric patients experiencing mild to severe AVH. Patients were recruited through health-care personnel in the Vestland County Health Care System (Helse Vest regional helseforetak). The recruitment was primarily from the Sandviken Psychiatric Clinic, Haukeland University Hospital in Bergen, Norway, with a few patients recruited from other counties. Patients had different psychiatric diagnoses, predominantly on the schizophrenia spectrum (see Supplementary Table 1 for ICD-10 diagnoses[13,14]). Prior to inclusion (and no more than 7 days before MR scanning), patients underwent a Positive and Negative Syndrome Scale (PANSS)[15] interview. Only patients exhibiting hallucinatory behaviour according to a PANSS positive subscale item 3 (P3) score of 3 or higher were recruited to the study. Subsequently, the project nurse administered the revised Beliefs About Voices Questionnaire (BAVQ-R)[16] and the MiniVoiceQuestionnaire (MVQ)[17]. Of the initially scanned participants, three patients were excluded due to technical issues, giving a total of 51 patients (19 female, 20 male, 2 transgender assigned female at birth), mean age 31.8 years (SD 9.3) with a mean PANSS P3 score of 4.6 (SD 0.8). Most patients used second-generation antipsychotics, with prescribed maximum dosage around 1.36 (SD 1.47) times the Defined Daily Dose (DDD)[18]; further detail in Supplementary Figure 1.

An equal number of healthy controls were included, matched on age (± 4 years): mean age 31.5 years (SD 9.1); two transgender patients were matched according to the sex they were assigned at birth. Healthy controls were drawn from a larger cohort which has been analysed previously to demonstrate efficacy of the applied methods [11].

All potential subjects were screened for implanted medical devices and history of major head injuries; controls were additionally screened for substance abuse, and for neurological or medical illnesses. All subjects provided written informed consent prior to participation and were free to withdraw at any time without consequence. Participants received compensation in the form of cash or a gift card. The study was approved by the Regional Committee for Medical Research Ethics in Western Norway (REK Vest # 2016/800).

#### 2.1.2 MR scanning protocol

MR data were acquired on a 3.0 T GE Discovery^TM^ MR750 scanner (GE HealthCare, Chicago, IL), with an 8-channel head coil. The protocol included a high-resolution T1- weighted structural acquisition: fast spoiled gradient (FSPGR) sequence with 188 sagittal slices of 256x256 isometric 1 mm voxels, 12 degree flip angle and TE/TR approximately 2.95/6.8 ms respectively. GABA-edited MRS data were acquired with a modified MEGA-PRESS sequence (TE=68 ms, TR=1500 ms, 15ms editing pulses at 1.9/7.46 ppm for edit-ON/-OFF respectively, simple 2-step phase cycle to minimize periodic confounds, 700 transients total), from a 22x36x23 mm (18.2 mL) voxel placed medially in the Anterior Cingulate Cortex (ACC). This voxel was centred on an imaginary line projected through the forward part of the pons, parallel with the brain stem as illustrated in Figure 1. The standard GE HealthCare MEGA-PRESS implementation had been modified to send per-TR trigger pulses for task synchronisation, and to periodically disable CHESS water suppression (every third transient) to allow acquisition of a water-unsuppressed reference signal interleaved within the regular GABA-editing sequence [11]. A summary of key sequence and hardware parameters is presented in an MRSinMRS[19] checklist, Supplementary Table 2.

**Figure 1:**
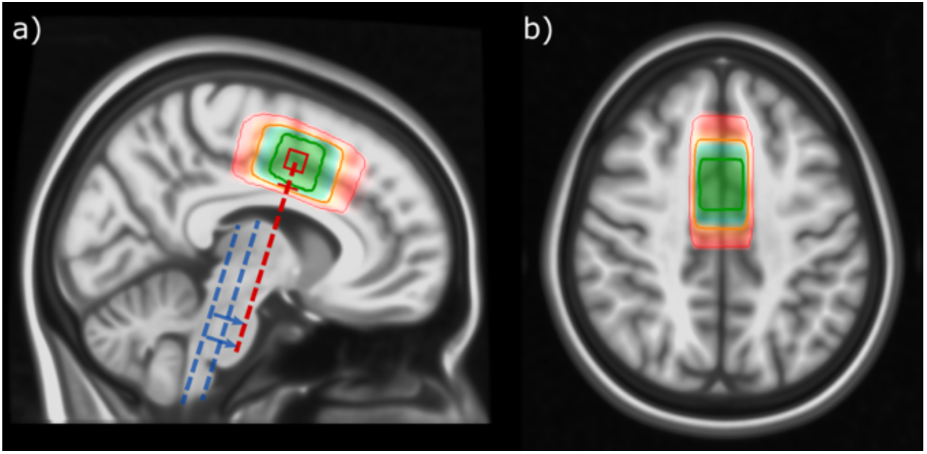
Placement of the fMRS voxel across all subjects, mapped to standard space. Shading (red-blue-green) and corresponding contours indicate [5,50,95]-percentile coverage of the achieved placement across subjects. Dashed lines in (a) illustrate landmarks used for voxel positioning: medial ACC, centred on an imaginary line through the forward part of the pons (red), parallel with the brain stem (indicated in blue). Adapted from Craven et al, 2023[11], updated to reflect the present sample.

Finally, BOLD fMRI data were collected with an echo-planar imaging (EPI) sequence: TE=30 ms, TR=2500 ms, 90 degree flip angle, interleaved acquisition of 36 slices of 128 x 128 voxels (1.72x1.72 mm), 3.0 mm slice thickness with 0.5 mm gap (3.5 mm slice spacing); 240 volumes for a total acquisition time of 600 s. Two patients elected to end scanning before completion of the BOLD fMRI task, and are therefore excluded only from analyses relating to that sequence.

#### 2.1.3 Functional paradigm: Eriksen Flanker task

During both the GABA-edited fMRS and the BOLD fMRI acquisitions, subjects performed a cognitive task based on the Eriksen Flanker task[20]. In each trial, a set of five arrows is presented; the task is to indicate the direction of the central “target” arrow. The four surrounding “flankers” may match the target for a “congruent” trial (for example, “ > > > > > ” or “ < < < < < ”), or they may be in the opposite direction for a cognitively more demanding “incongruent” trial (“ < < > < < ” or “ > > < > > ”). The paradigm was implemented in E-Prime 2.0 SP1 [2.0.10.353: Psychology Software Tools Inc., Pittsburgh, PA, https://pstnet.com/], with timing synchronised to the scanner via an NNL SyncBox [NordicNeuroLab AS (NNL), Bergen, Norway, http://nordicneurolab.com/, note declaration of interest]. Stimuli were presented through goggles (NNL) in light grey on a black background, with a small font to remain near the parafoveal field of view[21]. Subjects responded using handheld response grips (NNL), pressing with the index finger on the corresponding hand. Subjects were presented instructions in Norwegian or English according to preference, and were shown sample stimuli before entering the scanner, and again immediately before the task.

The task was implemented in a block-event design, beginning with a 60-second task-OFF block then alternating 30-second task-ON and 60-second task-OFF blocks, for a total of 11/6 task-ON blocks for the fMRS/fMRI acquisitions respectively. Within each task-ON block, one trail was presented per TR giving an average inter-stimulus interval (ISI) of 1500 ms and a total of 220/120 trials for fMRS/fMRI respectively. A randomly selected 40% of trials were incongruent. Trial onset timing was jittered randomly with respect to the fixed TR, such that stimuli were presented in the range T_S-A_ = 100-350 ms before the excitation pulse, spanning much of the early response suggested by prior fMRS studies[8,22]. Stimuli were presented for 350 ms, with a nominal 800 ms response window.

### 2.2 fMRS data processing and quantification

Functional MEGA-PRESS data were processed with a modified pipeline based around core functionality from Gannet version 3.1 [23]. After spectral registration[24,25], individual transients were processed using a linear model to separate variance of interest from nuisance factors (variance associated with phase cycling and inferred subject motion), as detailed in a previous work [11,26]. Spectra were modelled according to achieved interval between stimulus and acquisition (T_S-A_). Five bins were defined, with edges at [100, 183, 267, 350] ms, open at either end, lower limits inclusive; the inner three bins evenly cover the nominated 100-350 ms T_S-A_ range. This resulted in approximately 48 task-ON metabolite transients per bin (with some individual variation), and 320 task-OFF metabolite transients. Line-shape matching was performed using a reference deconvolution approach[26–29].

Extracted GABA-edited difference spectra (DIFF) were evaluated against three rejection criteria, applied in series: full width at half maximum (FWHM) linewidth of fitted GABA+ or inverted N-acetylaspartate (NAA) peaks exceeding 30 Hz or 12 Hz respectively, extraordinarily low signal-to-noise ratio (SNR) (below 20 for the fitted NAA peak), and extreme outliers, where GABA+ or Glx estimates differed from the median by more than five times the Median Absolute Deviation (MAD) for spectra surviving the first two criteria.

Unsuppressed water-reference spectra (WREF) were fit with a pseudo-Voigt function on a linear baseline, with Voigt linewidth filtered for outliers and discontinuities and modelled against the expected BOLD response (event impulses convolved with a dual-gamma haemodynamic response function (HRF) model), with covariate components as for the metabolite spectrum model. The resultant BOLD model coefficient estimates the overall change in water linewidth (ΔFWHM_water_) reflecting the strength of the individual’s BOLD response within the fMRS-localised region[11]. This value will be denoted BOLD-fMRS.

### 2.3 fMRI data processing

fMRI block analysis was performed using FEAT (FMRI Expert Analysis Tool version 6.00, part of FSL) [30–37], with a standard pipeline described fully in our previous work[11]. For the present study, a per-subject volume of interest (VOI) is defined from the individually prescribed fMRS voxel geometry in the ACC (VOI_fMRS,ACC_), with the median Z-score (without thresholding) across VOI_fMRS,ACC_ taken as a measure of the strength of the BOLD response across that volume. This value will be denoted BOLD-fMRI.

### 2.4 Numerical and Statistical Analysis

Behavioural outcomes were assessed with the Wilcoxon Signed Rank test for related samples (between session/stimulus, within subject), and the Mann-Whitney U-test for unrelated samples (patient vs control), motivated by unequal variance and high skew in some parameters. Correlation of BOLD estimates by the two methods (BOLD-fMRI, BOLD-fMRS) was assessed using the skipped Spearman method[38,39] for resilience to bivariate outliers.

Outcomes from hypothesis testing and correlational tests are adjusted for multiple comparisons within sub-analysis, using the Holm-Bonferroni approach; adjusted p-values are denoted p_holm_, with a corrected significance threshold defined as p_holm_<0.05.

Least-squares linear modelling was used to assess associations between baseline metabolite estimates, BOLD signal strength and interactions with patient and control groups, with voxel grey matter fraction (fGM) as a covariate (ie, `Glx ∼ C(group)*BOLD + fGM`). Outlier observations having disproportionate influence on the model (according to the studentized difference in fits[40] thresholded at 2√(k/n)) were dropped. Model suitability was verified using the Jarque-Bera test of normality[41], and White’s Lagrange Multiplier and Two-Moment Specification tests[42] for heteroscedasticity and correct specification.

A series of exploratory correlational tests were performed between symptom scores (PANSS: P3, total positive and total negative subscale scores) and baseline metabolite concentrations (Glx, GABA+), task-elicited change in metabolite levels (ie, ΔGlx = Glx_task-ON_ - Glx_task-OFF_ and ΔGABA = GABA_task-ON_ - GABA_task-OFF_), BOLD response, and task performance metrics (response accuracy (RA)/reaction time (RT), reaction time slowing (RT_slowing_)), using the skipped Spearman method.

Finally, a linear mixed-effects model was constructed for metabolite estimate in relation to group (patient, control), task status (task-OFF, task-ON), with grey matter fraction as a covariate and subject as the grouping variable (ie, Glx ∼ C(group)*C(task_state) + fGM), filtering observations with strong residuals (deviating from median residual by more than 2.5 times the MAD).

Statistical testing was performed in Python (v3.9.17) with statsmodels [43] (v0.13.5), pingouin [44] (v0.5.2), SciPy [45] (v1.9.3), pandas [46] (v1.5.2) and NumPy [47] (v1.23.5) libraries; subsequent visualisation was built on tools from matplotlib [48] (v3.3.4), seaborn [49] (v0.11.2) and statannotations [50] (v0.5).

## 3 Results

### 3.1 Behavioural Outcomes

Behavioural outcomes from the Flanker task are summarised in Figure 2 and Supplementary Table 3. Task performance in terms of reaction time (RT), response accuracy (RA) and RA/RT was significantly degraded (p_holm_<0.001) between congruent and incongruent trials, assessed across all subjects. Improved RA and RA/RT were observed between the fMRS and subsequent fMRI task, both strongly significant for incongruent stimuli (p_holm_<0.001). Patients showed significantly lower RA and RA/RT than the healthy controls (p_holm_<0.001).

**Figure 2.**
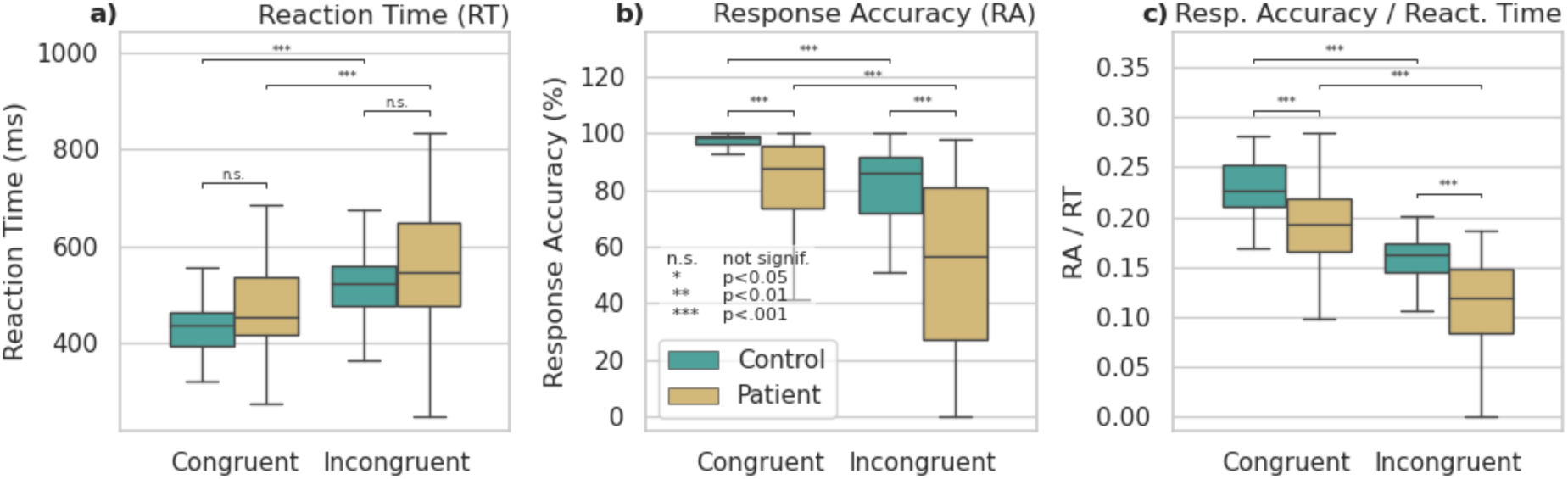
Outcomes from the behavioural task (fMRS and fMRI sessions pooled), showing significant increase in reaction time (a) and reduction in response accuracy (b, c) for incongruent conditions, and significantly degraded response accuracy for patients. Significant differences indicated with *** p_holm_<0.001, ** p_holm_<0.01, * p_holm_<0.05, n.s. not significant

### 3.2 Functional Outcomes

#### 3.2.1 BOLD assessment by fMRI and fMRS

Correlation between the strength of the BOLD response as assessed with fMRS and fMRI methods within the individually prescribed VOI_fMRS,ACC_ was significant for healthy controls (r=0.35, 95% confidence interval (CI_95%_) [0.07,0.57], p=0.014), but unreliable for the patient group (r=-0.08, CI_95%_ [-0.353,0.212], p>0.5), combining to a weak correlation across the entire dataset (r=0.16, CI_95%_ [-0.04,0.346], n.s.). Correlation by group is shown in Figure 3; note generally weaker BOLD-fMRI response and greater variance in the BOLD-fMRS estimate amongst patients.

**Figure 3.**
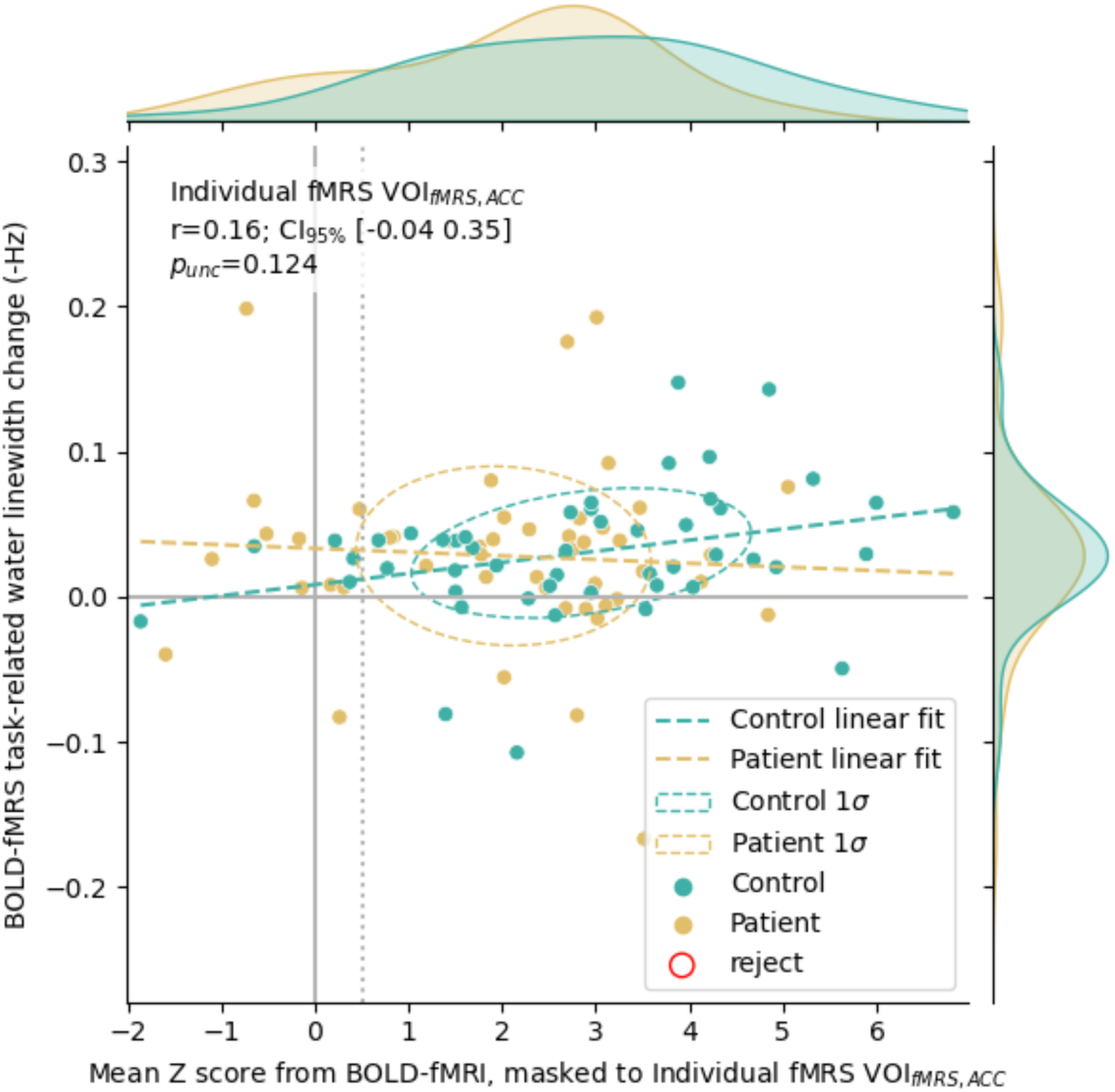
Relation of BOLD assessed by BOLD-fMRS linewidth changes (ΔFWHM_water_) to mean Z score observed from the BOLD-fMRI data, regionally masked to the individual fMRS voxel (VOI_fMRS,ACC_)), showing significant correlation specific to healthy controls.

#### 3.2.2 Baseline and Functional MRS

Mean measured spectra for each group and condition are presented in Figure 4, with corresponding quality metrics in Supplementary Table 4. Association with BOLD-fMRI is shown in Figure 5. The linear model for baseline metabolite concentration with factors for the interaction between group and measured BOLD strength (BOLD-fMRI) and grey matter fraction showed significant effects of group for Glx (-2.3603 i.u., CI_95%_ [-3.680, -1.040], p<0.001), a main effect for BOLD-fMRI (b=-0.394, CI_95%_ [-0.686,-0.102], p<0.01) and an interaction between group and BOLD-fMRI (b=0.586, CI_95%_ [0.113, 1.059], p=0.016). This indicated lower baseline Glx in patients and differential association between BOLD and baseline Glx amongst healthy controls (negative association) and patients (positive association). All other factors were non-significant. Substituting BOLD-fMRS as the independent variable yielded weaker outcomes (p=0.054 for the group effect, p=0.094 for the main effect of BOLD-fMRS). Similar analyses for GABA+ yielded no significant results.

**Figure 4.**
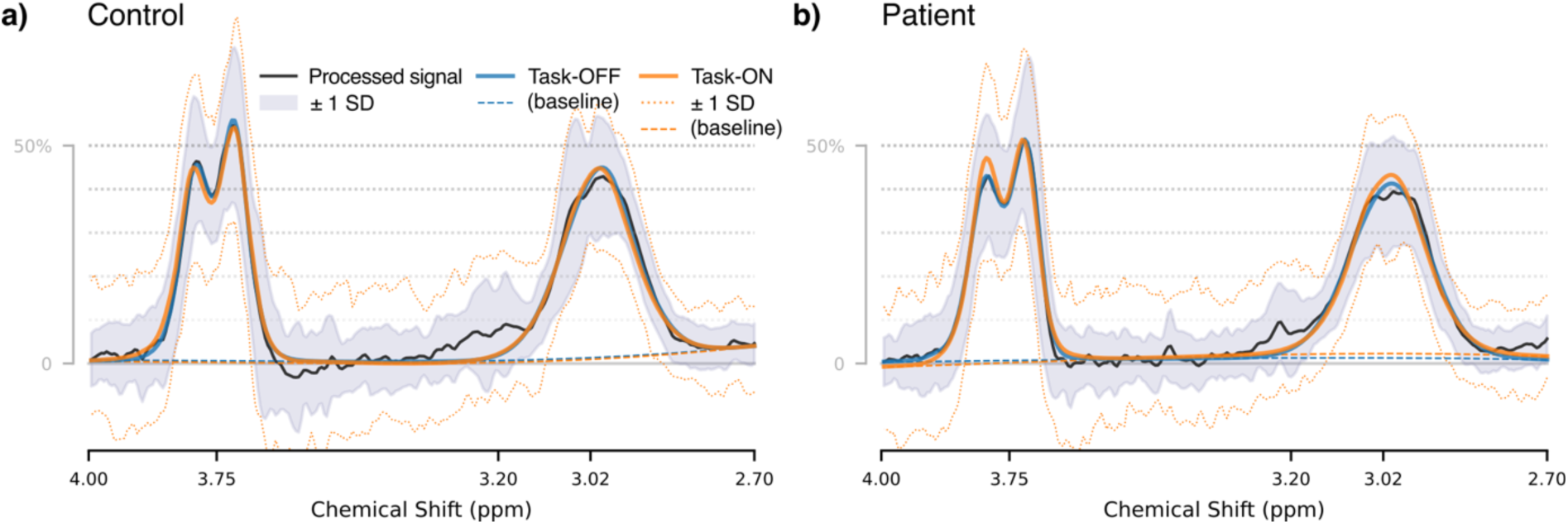
Group mean spectra and model fit, separated by task condition

**Figure 5.**
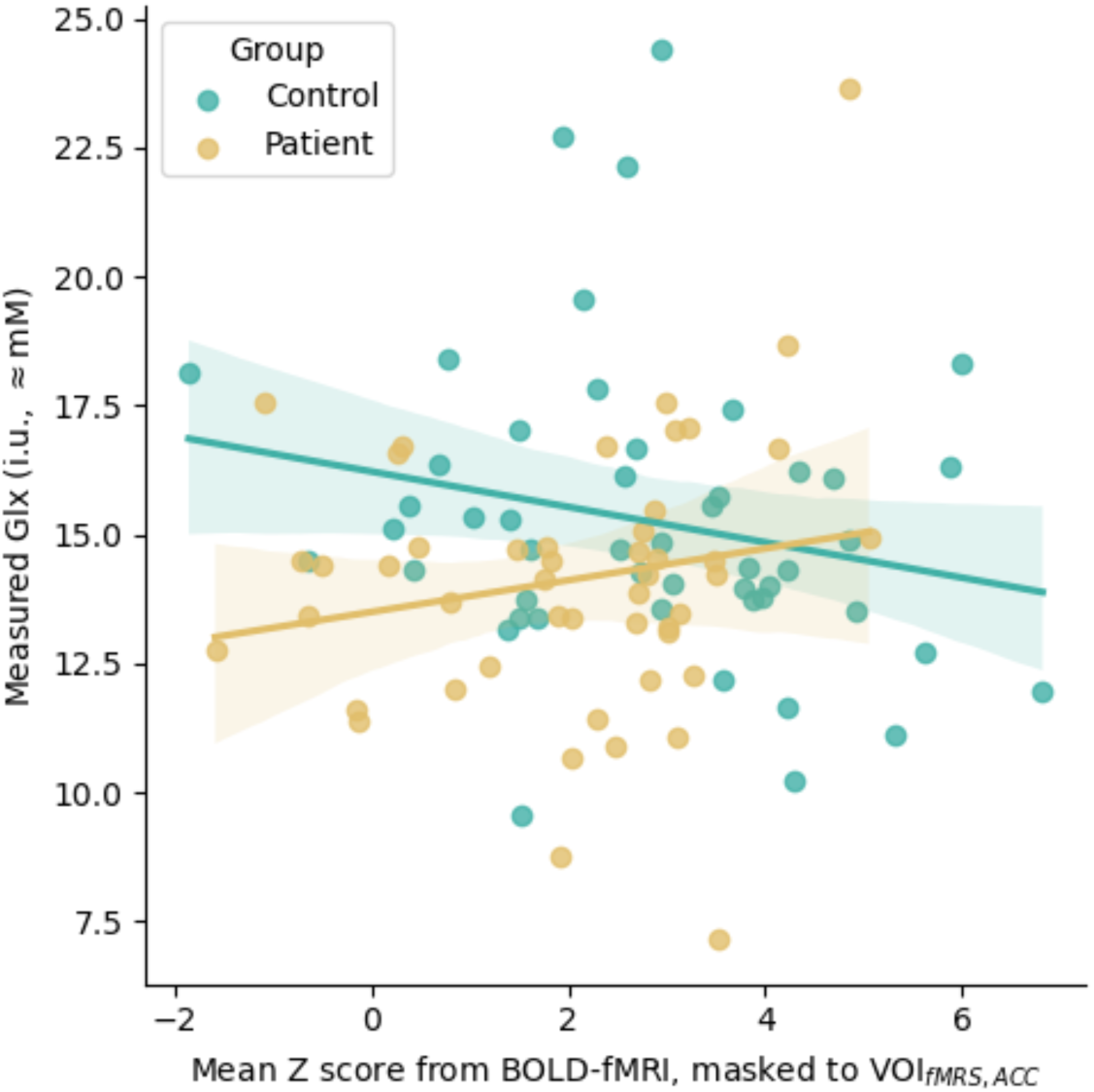
Relation between measured BOLD-fMRI response and measured Glx, showing complementary effects for healthy controls and patients.

Detailed model output is presented in the Supplementary Material, sections C.1 and C.2 for Glx and GABA+ respectively.

No correlation was seen between symptom scores (PANSS P3, total positive, total negative) and baseline metabolite concentration (GABA+ or Glx) or BOLD response (all p_holm_>0.05). Possible associations between behavioural reaction time slowing and total negative symptoms (r=-0.351, CI_95%_ [-0.578, -0.0747], p=0.0143) and between total positive symptoms and ΔGlx (r=-0.335, CI_95%_ [-0.57, -0.0496], p=0.0228) did not survive strict correction for multiple comparisons (p_holm_=0.3, 0.46 respectively) in the present context.

Outcomes for all exploratory tests are presented in Supplementary Material, section C.3 (Supplementary Table 5).

Concentration estimates for GABA+ and Glx by group and task condition are presented in Figure 6; additional metabolites estimated from the edit-OFF sub-spectrum (Choline, Creatine, NAA) are presented for interest in Supplementary Figure 2, but not investigated further. Mixed-effects linear modelling showed strong task-related increase in Glx (1.107 i.u., CI_95%_ [0.343 1.871], p<0.01) against an intercept at 13.096 i.u., CI_95%_ [7.737,18.455], or roughly 8.5% increase in Glx estimate in response to the functional task. The same model indicated reduced baseline Glx in the patient groups (-1.08 i.u., CI_95%_ [- 2.130, -0.030], p=0.04), but no evidence of any interaction was found (i.e., no differential effects between task-ON and task-OFF, for patients vs controls (p>0.5)). Similar modelling for GABA+ showed no significant effects.

**Figure 6.**
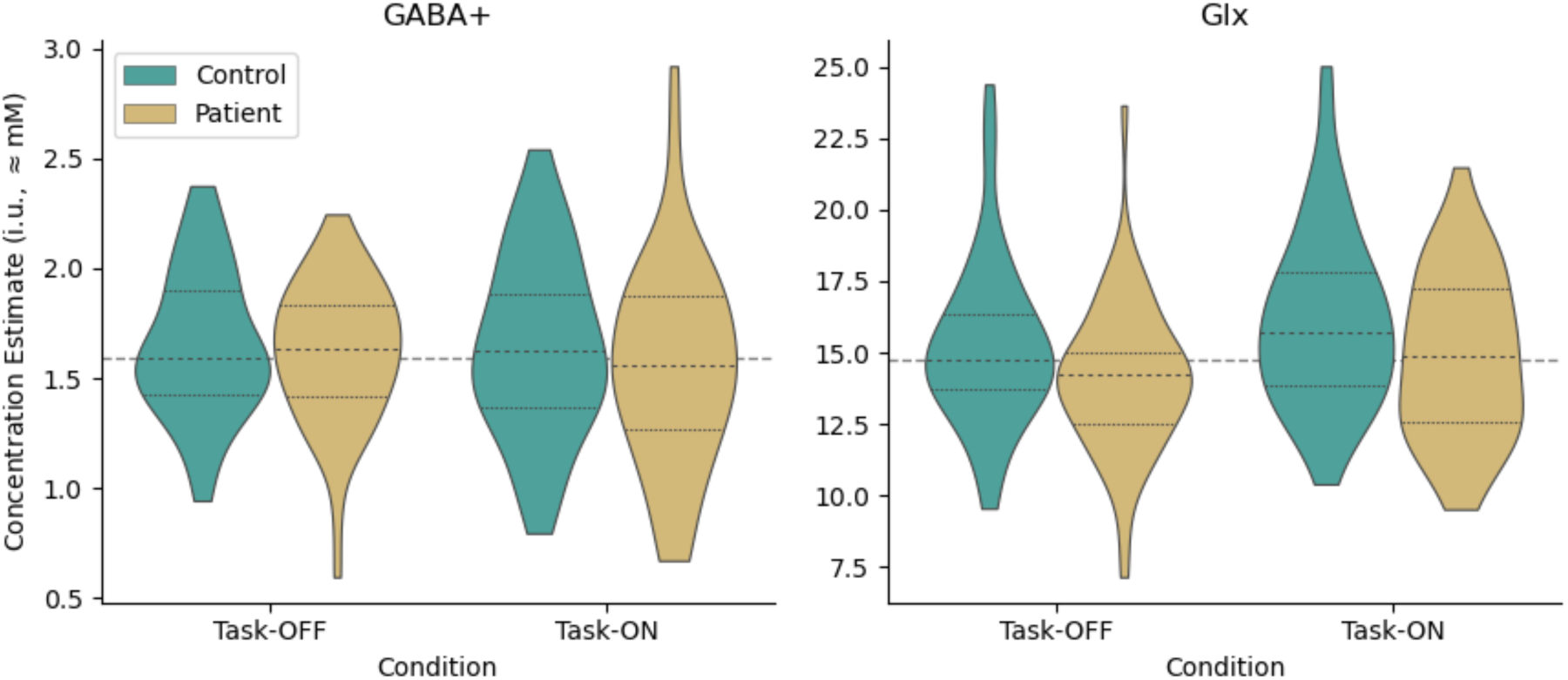
Metabolite estimates by task state and group

## 4 Discussion and Conclusions

### 4.1 Behavioural outcomes

Behavioural outcomes were consistent with expectations based on existing literature[21,51–54], including substantially reduced response accuracy in the patient group indicative of an executive and attentional deficit. However, anticipated prolonged reaction times[55,56] in the patient group were not significant, likely due to high variance within that group. A previously reported improvement in response accuracy between the fMRS and subsequent fMRI task performance[11], perhaps attributable to learning effects, was also apparent within the current patient group.

### 4.2 Association with Symptom Scores

Lack of observed correlation between symptom scores (PANSS P3, total positive, total negative) and either the strength of the BOLD response or baseline metabolite concentration may at first seem surprising, especially given previous reports showing such correlations[57–60] (albeit with some contradictory findings[61]). However, we note that high PANSS P3 was a pre-condition for inclusion in the present study, meaning the distribution of symptom scores is not fully representative of the general population of patients with psychosis. Therefore, the absence of correlations in the present study is not necessarily in conflict with existing reports; it may be a result of sensitivity/ceiling effects. Moreover, it may indicate that these factors distinguish low/non-hallucinating subjects from high-hallucinating ones, rather than degrees of severity amongst the higher-hallucinating patients.

### 4.3 Methodological Considerations

Our previous study on healthy controls showed a significant correlation between the BOLD signal strength assessed by the fMRS and subsequent fMRI acquisition. The moderate degree of this correlation was attributed to inherent variability of each measure, intra-session variability in the BOLD response itself, possible learning effects and/or fatigue. No such correlation could be demonstrated in the patient group of the present study; there are a few possible explanations for this. Within the patient group, BOLD-fMRS measurement showed greater variance than amongst controls – perhaps reflecting reduced reliability of the measurement in the presence of more pronounced subject motion as typically seen amongst patient groups. BOLD estimates for patients also exhibited a skew towards lower values, particularly evident in the BOLD-fMRS values (Figure 3) and likely reflecting limited conformance amongst some of the participants. Weaker BOLD effects may challenge the sensitivity of the fMRS implementation.

Hence, while the BOLD-fMRS estimates obtained in this way remain useful in certain scenarios (especially where a robust BOLD response is to be expected) and offer the great advantage of temporal concurrency with the MRS acquisition, we would caution against relying solely upon this technique in scenarios where the BOLD response may be more subtle or more variable, or where subject conformance may be limited. Further refinement of the analysis model could potentially improve performance in this regard.

### 4.4 Baseline Metabolite Concentrations

Our results indicated lower baseline Glx in patients relative to controls in the ACC region. While this is compatible with some previous reports for the region[60–62], recent meta- analyses [63–65] have found differing effects depending on the brain region, and more nuanced associations have been reported in relation to age[66], progression/chronicity[67,68], antipsychotic effects[69] and treatment response[65,70,71]. Studies have also shown correlation between glutamate level and grey matter loss in schizophrenia[72], consistent with neurodegeneration (loss of cortical volume, presumably corresponding with a decrease in glutamatergic synaptic density[73]) and likely associated with age and duration of illness. Inclusion of the fGM factor in our statistical models should limit the impact of this factor on our present findings.

Interpretation of our Glx findings is constrained by a few technical limitations: the meta-analyses suggest different effects for Glu and Gln when assessed separately, with ACC Gln tending to show elevation in patients while ACC Glu exhibits decreases in particular sub-groups. This may reflect reduced glutamine demand, perhaps as a result of N-methyl-ᴅ- aspartic acid (NMDA) receptor hypofunction[74], leading to a net accumulation of glutamine[73]. While no attempt was made to distinguish Glu from Gln in the current dataset, the DIFF spectrum of the TE=68 ms sequence is likely to differ from un-edited, shorter TE sequences in terms of relative sensitivity to each. Inability to distinguish synaptic glutamate from other roles (energy metabolism, protein synthesis etc) may also limit interpretation of these outcomes. Furthermore, our selection criteria target a specific symptom (hallucinatory behaviour) rather than a particular diagnosis. This yields a sample which is heterogeneous with respect to diagnosis, treatment strategy and (presumably) genetic factors; finer-grained analysis considering some of these factors presents an opportunity for further investigation.

We also observe a differential relation between baseline ACC Glx and BOLD-fMRI in the same region: a negative association in healthy controls, contrasting a positive association in patients. This is consistent with the findings of Falkenberg et al [12], and compatible with altered glutamatergic function in schizophrenia[75–78]. Indeed these findings may reflect an underlying neuronal mechanism, with positive association in the patient group perhaps compensatory for overall reduced BOLD activation, with increasing glutamate levels reflecting higher rates of energy turnover in the region [79,80].

### 4.5 Dynamic Metabolite Concentrations

The finding of increased Glx in response to task (approximately 8.4% increase between task-OFF and task-ON states, across groups) is consistent with our previous study[11] which investigated healthy controls alone, and towards the upper edge of ranges reported in previous meta-analyses (for example, 6.97% CI_95%_ [5.23, 8.72] change reported by Mullins [81], although typically lower (∼3%) for the few studies investigating basic cognitive tasks in the ACC [82–84]). A short-term dynamic change of this magnitude is unlikely to be explained by metabolic processes alone, but may be consistent with a hypothesized compartmental shift[81,85]. During neural activity, Glu may move from a pool with reduced MRS visibility [86–88] (such as the presynaptic vesicle) to a compartment where it is more visible (such as the cytosol or the synapse), leading to an increase in the MRS-measured signal.

Significantly, the relative task-related change in Glx levels did not differ between healthy controls and patients. Combined with the baseline outcomes, this suggests that existing findings of glutamatergic effects are driven by lower baseline Glx levels in patients, rather than aberrant dynamic regulation in response to task. Furthermore, the absence of observable effects for GABA, either at baseline or as a change in response to task, suggests that neurometabolic effects underlying a putative excitatory/inhibitory imbalance are again driven by baseline Glx deficit, rather than GABAergic abnormalities. Although broadly compatible with the notion of an excitatory-inhibitory imbalance, these findings are less clear in relation to putative NMDA receptor hypofunction[74] and its consequences for GABAergic functions[89]; indeed, it has been proposed that NMDA receptor hypofunction would lead to increased release of synaptic glutamate and underdeveloped GABAergic circuitry.

### 4.6 Conclusion

In summary: with concurrent measurement of GABA+, Glx and BOLD in response to a cognitive task, we obtained behavioural, functional MR and spectroscopy findings consistent with literature based on separate acquisition of fMRI and MRS, and indicative of an executive and attentional deficit. Limited findings in relation to symptom severity are likely due to limited variation in our defined sample. Reduced baseline Glx in patients, together with a dynamic change comparable with healthy controls in response to task points to tonic rather than phasic effects. Together with an absence of observable differences for GABA this serves to refine our understanding of the roles of these metabolites in a putative

Excitatory/Inhibitory imbalance as a possible mechanism underlying the perception of auditory hallucinations.

## 5 Author Contributions

ARC: Conceptualization, Methodology, Software, Validation, Formal analysis, Investigation, Data Curation, Writing – Original Draft

GD: Conceptualization, Methodology, Validation, Investigation, Writing – Review & Editing LE: Methodology, Validation, Resources, Writing – Review & Editing

KK: Data Curation, Writing – Review & Editing

LL: Investigation, Data Curation, Writing – Review & Editing RN: Methodology, Software, Writing – Review & Editing LBS: Investigation, Data Curation, Writing – Review & Editing

EJ: Conceptualization, Resources, Writing – Review & Editing, Project administration

KH Conceptualization, Methodology, Resources, Writing – Review & Editing, Supervision, Project administration, Funding acquisition

## Acknowledgements

This study was funded by the European Research Council (ERC) grant #693124 and by the Western Norway Health Authorities (Helse-Vest) grant #912045 to Kenneth Hugdahl. We are grateful for the radiographers at Haukeland University Hospital: Roger Barndon, Christel Jansen, Turid Randa, Trond Øveraas, Eva Øksnes and Tor Erlend Fjørtoft, for their time and patience with data collection throughout this study.

## 6 Declaration of interest

Co-authors ARC, LE, KH own shares in NordicNeuroLab (NNL), which produced some of the hardware accessories used during functional MR data acquisition at the scanner. The authors declare no other conflicting interests.

## 7 Data availability statement

In accordance with data sharing regulations imposed by the Western Norway Ethical Committee (REK-Vest) (https://rekportalen.no/), data may be shared by request to the corresponding author, subject to written permission from the REK-Vest.

Our custom MEGA-PRESS sequence is based on proprietary GE HealthCare code; the authors are in principle willing to share details on our local adaptions through the appropriate vendor-facilitated channels.

Our tools and pipelines for fMRS data modelling are available from: https://git.app.uib.no/bergen-fmri/gaba-temporal-variability

## List of Abbreviations

(f)MRI: (functional) Magnetic Resonance Imaging
(f)MRS: (functional) Magnetic Resonance Spectroscopy
ACC: Anterior Cingulate Cortex
AVH: Auditory Verbal Hallucination
BAVQ-R: revised Beliefs About Voices Questionnaire
BOLD: blood-oxygenation level dependent
CI_95%_: 95% confidence interval
DDD: Defined Daily Dose
DIFF: GABA-edited difference spectrum
EPI: echo-planar imaging
fGM: voxel grey matter fraction
FWHM: full width at half maximum (measuring spectral linewidth)
GABA: γ-aminobutyric acid
GABA+: GABA with contribution from underlying co-edited signals
Glu: glutamate
Glx: Glutamate and Glutamine combined
HRF: haemodynamic response function
ISI: inter-stimulus interval
MAD: Median Absolute Deviation
MVQ: MiniVoiceQuestionnaire
NMDA: N-methyl-ᴅ-aspartic acid
NAA: N-acetylaspartate
P3: PANSS positive subscale item 3, hallucinatory behaviour
PANSS: Positive and Negative Syndrome Scale
RA: response accuracy
RT: reaction time
RT_slowing_: reaction time slowing
SNR: signal-to-noise ratio
SSD: schizophrenia spectrum disorder
T_S-A_: Time from Stimulus to Acquisition
VOI_(location)_: volume of interest (in specified location)
WREF: Unsuppressed water-reference spectrum

## A Additional Subject Details

**Supplementary Table 1.**
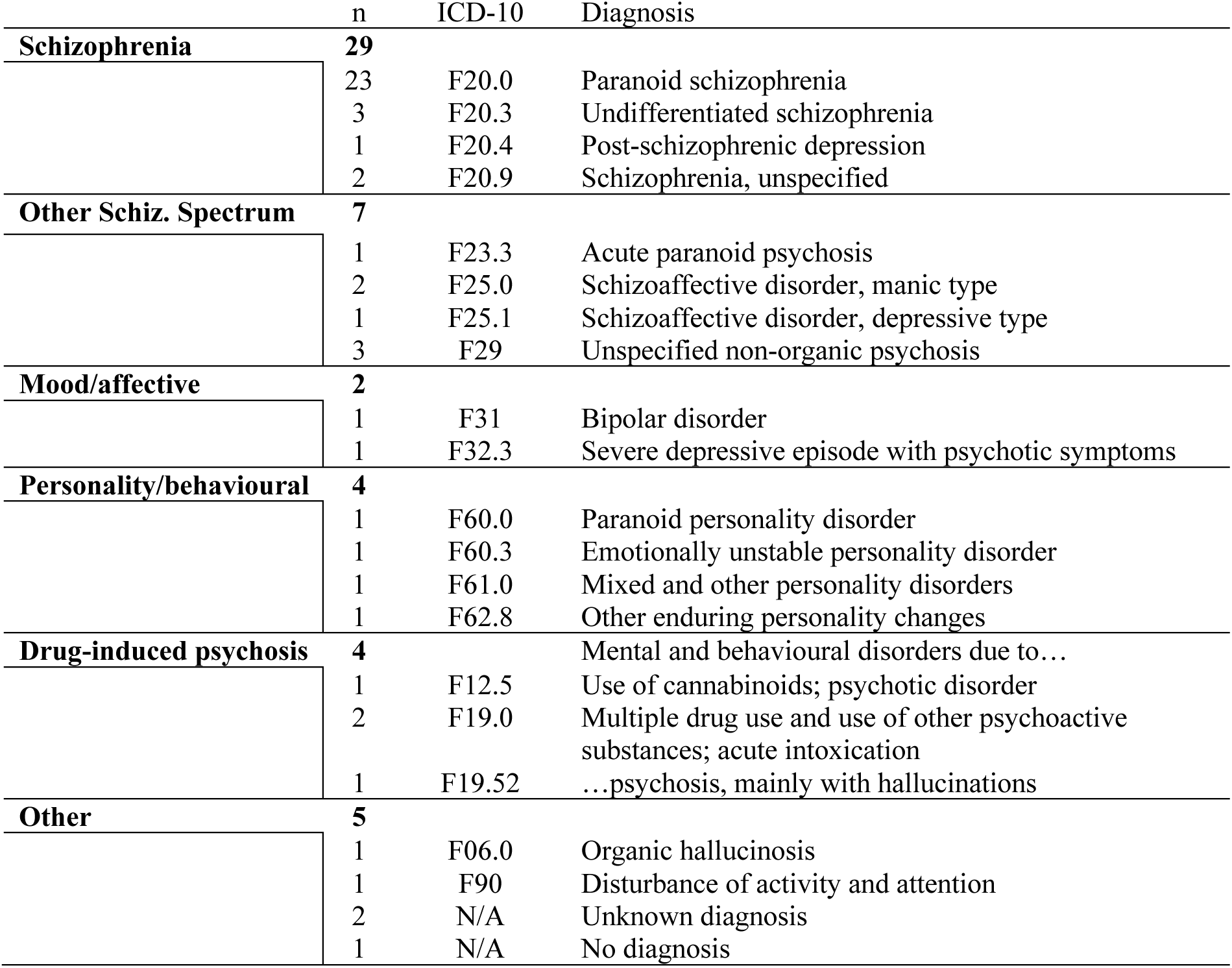
Diagnoses of patients in the present study, according to ICD-10 criteria.

Diagnoses are according to the ICD-10 criteria, described in the following publications (English and Norwegian translation):

World Health Organization. (1992). *The ICD-10 classification of mental and behavioural disorders: Clinical descriptions and diagnostic guidelines* (Reprinted). World Health Organization. https://www.who.int/docs/default-source/classification/other-classifications/bluebook.pdf

World Health Organization. (2016). *ICD-10 psykiske lidelser og atferdsforstyrrelser: Kliniske beskrivelser og diagnostiske retningslinjer* (10. rev., 19. oppl). Universitetsforlaget. https://www.ehelse.no/kodeverk-og-terminologi/ICD-10-og-ICD-11

**Supplementary Figure 1.**
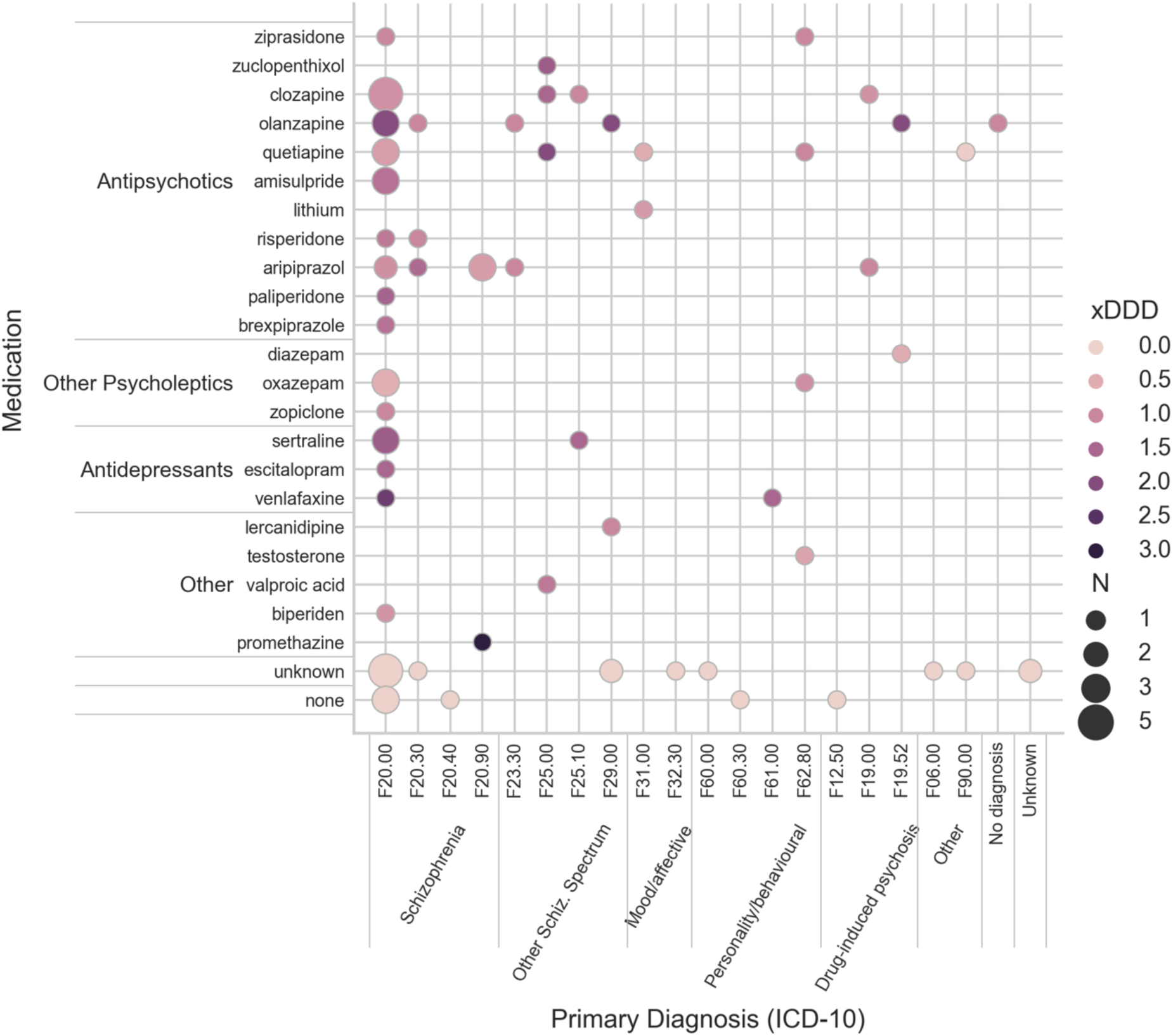
Medication and mean of the (maximum) prescribed dosage, expressed relative to the defined daily dose (DDD). Size of the points is proportional to the number of patients (N), with darker shading indicating higher dosage (xDDD).

In several cases, the prescribed dosage was qualified with terms such as “up to” and “as needed” (“opp til”, “ved behov”); some subjects reported that they had not taken prescribed medicine in the period leading up to the study. Dosage presented here therefore represents the maximum prescribed dosage, and is likely to slightly overestimate the actual dosage taken.

Defined daily dose (DDD) according to WHO Collaborating Centre for Drug Statistics Methodology, ATC classification index with DDDs, 2024. Oslo, Norway 2024, searchable online at https://atcddd.fhi.no/atc_ddd_index/

ATC codes were derived from trade names via the register at https://www.felleskatalogen.no/medisin/atc-register/

## B MRSinMRS checklist

**Supplementary Table 2.**
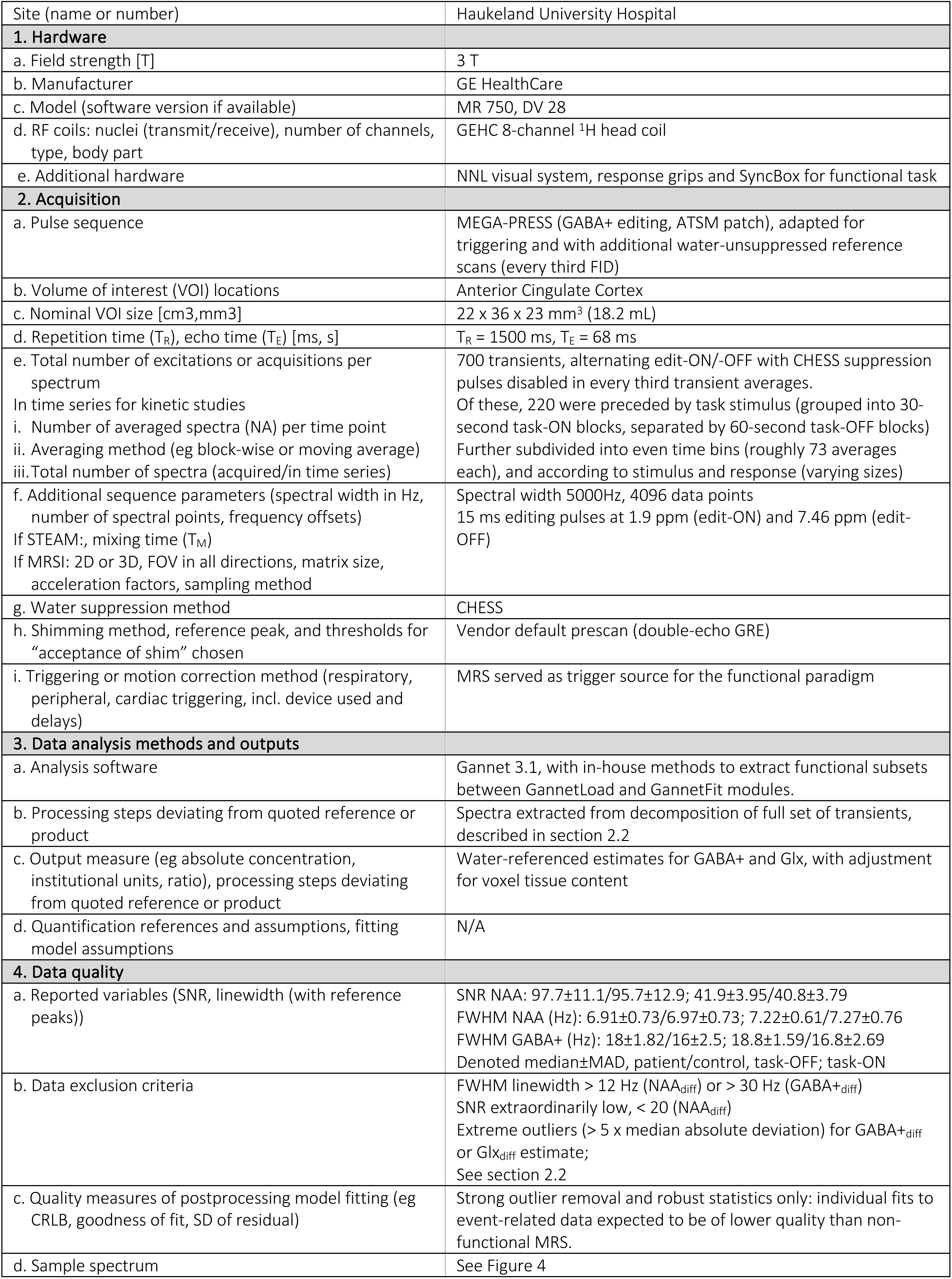
MRSinMRS checklist[19] summarising key details of the MRS acquisition.

## C Supplementary Results

**Supplementary Table 3.**
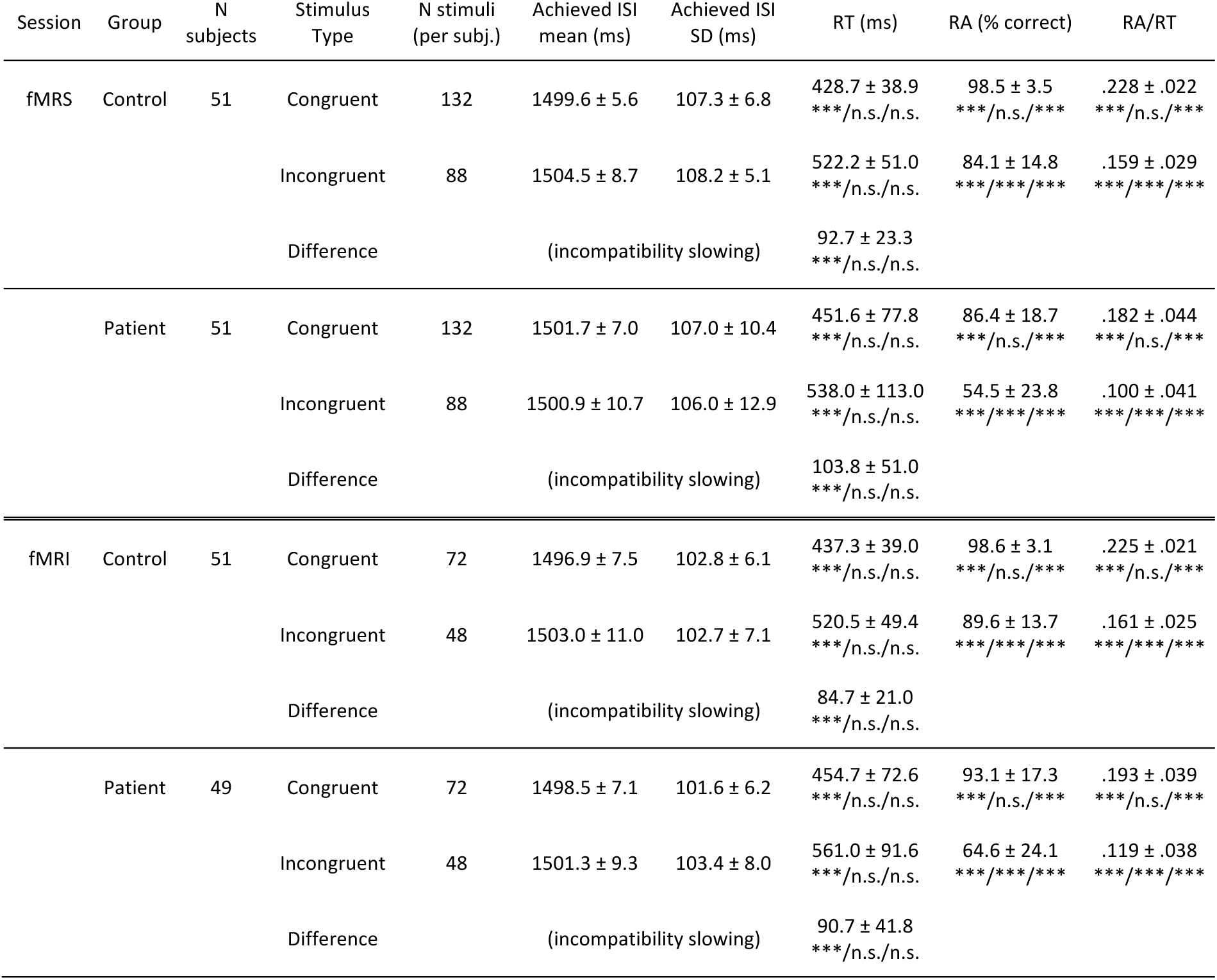
Behavioural outcomes from the Flanker task; values are quoted as Median +/- Median Absolute Deviation (MAD) of per-subject outcomes. Significant differences are indicated between stimulus type, session and group (denoted type/session/group, *** p_holm_<0.001, ** p_holm_<0.01, * p_holm_<0.05, n.s. not significant). ISI: Inter-stimulus interval, RA: Response Accuracy, RT: Response Time.

**Supplementary Table 4.** Quality metrics and concentration estimates from the fMRS analysis, task-ON vs task-OFF, presented as median ± MAD.

|  | Control |  | Patient |  |
| --- | --- | --- | --- | --- |
|  | task-OFF | task-ON | task-OFF | task-ON |
| SNR NAA | 97.7 ± 11.1 | 41.9 ± 3.95 | 95.7 ± 12.9 | 40.8 ± 3.79 |
| FWHM Cho/Cr *before* linewidth matching n.s. | 6.84 ± 0.569 | 6.72 ± 0.619 | 7.07 ± 0.474 | 7.02 ± 0.518 |
| FWHM Cho/Cr *after* linewidth matching n.s. | 6.88 ± 0.569 | 7.03 ± 0.651 | 7.24 ± 0.54 | 7.23 ± 0.509 |
| FWHM NAA (Hz) | 6.91 ± 0.729 | 6.97 ± 0.729 | 7.22 ± 0.608 | 7.27 ± 0.764 |
| FWHM GABA+ (Hz) | 18 ± 1.82 | 16.3 ± 2.5 | 18.8 ± 1.59 | 16.8 ± 2.69 |
| FWHM Glx (Hz) | 12.1 ± 1.11 | 12.7 ± 1.47 | 12.3 ± 0.9 | 12.5 ± 1.49 |
| GABA+ (i.u., ≈mM) | 2.9 ± 0.367 | 2.88 ± 0.46 | 2.87 ± 0.372 | 2.79 ± 0.594 |
| Glx (i.u., ≈mM) | 14.7 ± 1.35 | 15.7 ± 2.03 | 14.2 ± 1.33 | 14.9 ± 2.39 |

**Supplementary Figure 2.**
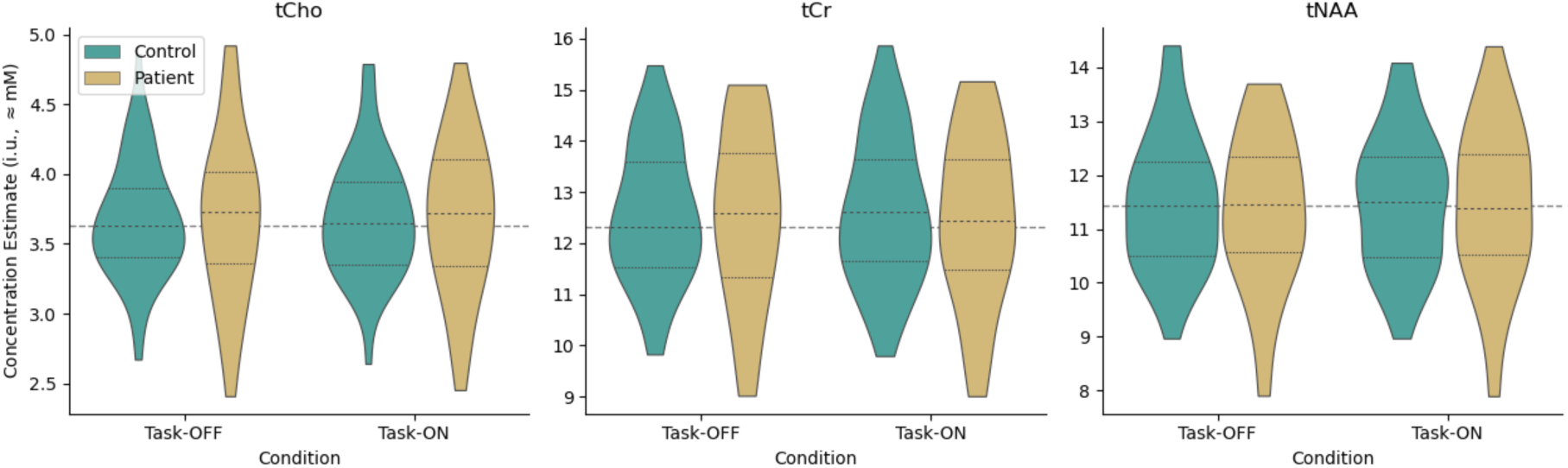
Concentration estimates for other metabolites according to group and condition (obtained from the edit-OFF sub-spectrum using the Gannet peak-fitting model)

### C.1 Regression Modelling Outcomes: Glx

Associations between baseline Glx estimate, BOLD signal strength and interactions with patient and control groups, with voxel grey matter fraction fGM as a covariate (ie, Glx ∼ C(group)*BOLD + fGM), after removing outlier observations:

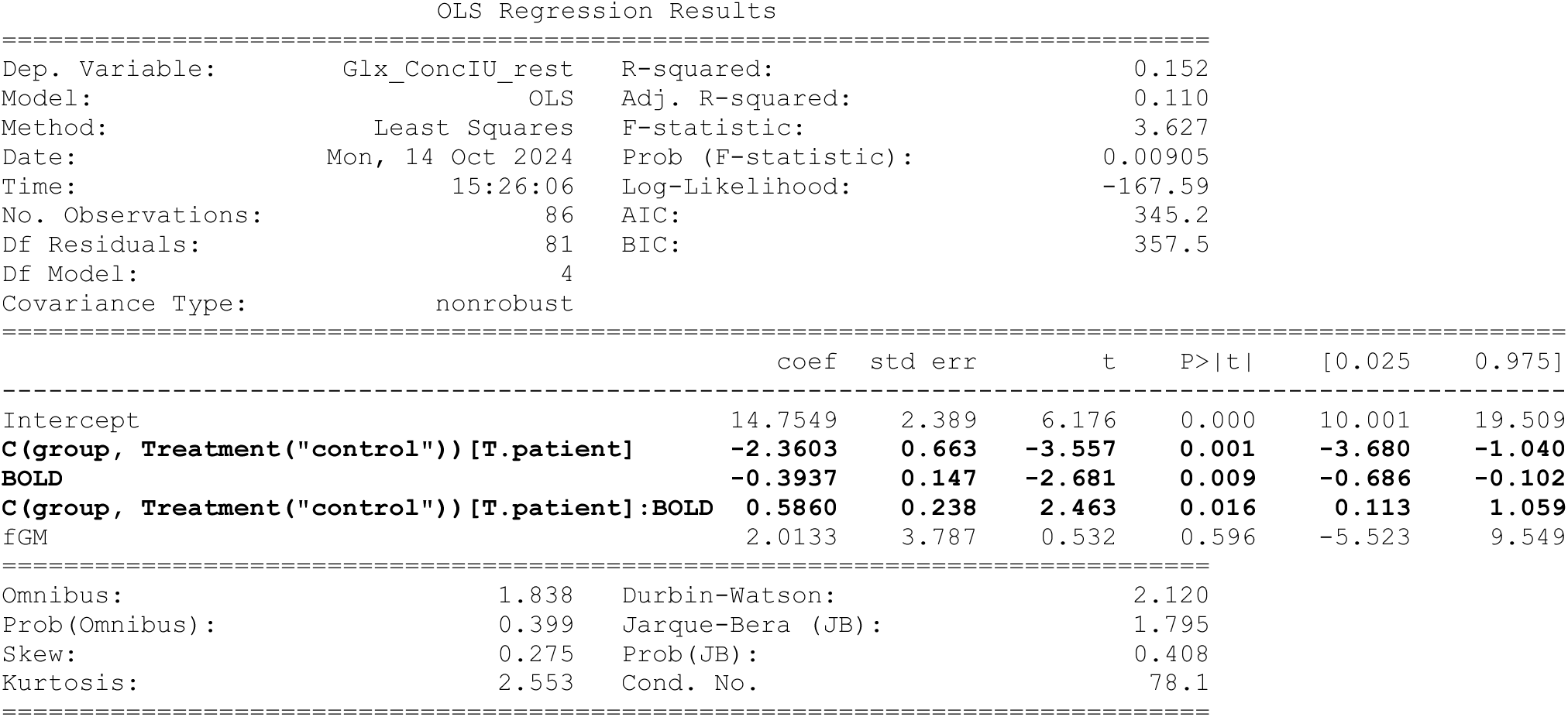

Glx estimate in relation to group (patient, control), task status (task-OFF, task-ON), with grey matter fraction as a covariate and subject as the grouping variable (ie, Glx ∼ C(group)*C(task_state) + fGM), after filtering observations with strong residuals:

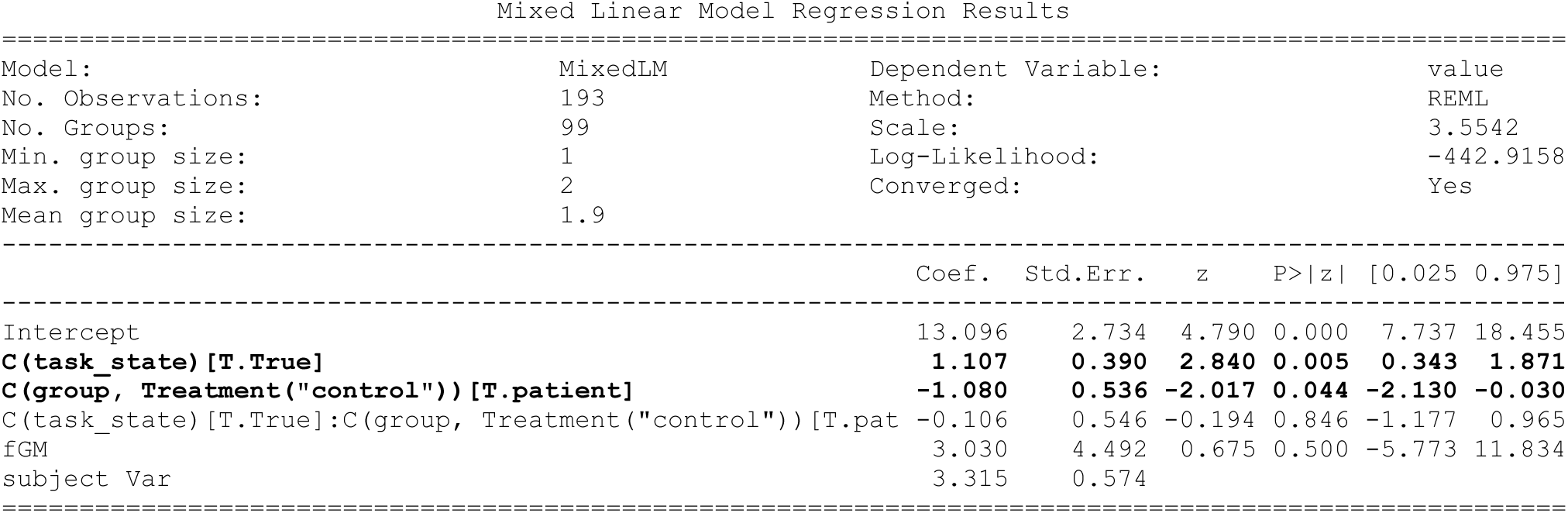

### C.2 Regression Modelling Outcomes: GABA+

Associations between baseline GABA+ estimate, BOLD signal strength and interactions with patient and control groups, with voxel grey matter fraction fGM as a covariate (ie, GABA ∼ C(group)*BOLD + fGM), after removing outlier observations:

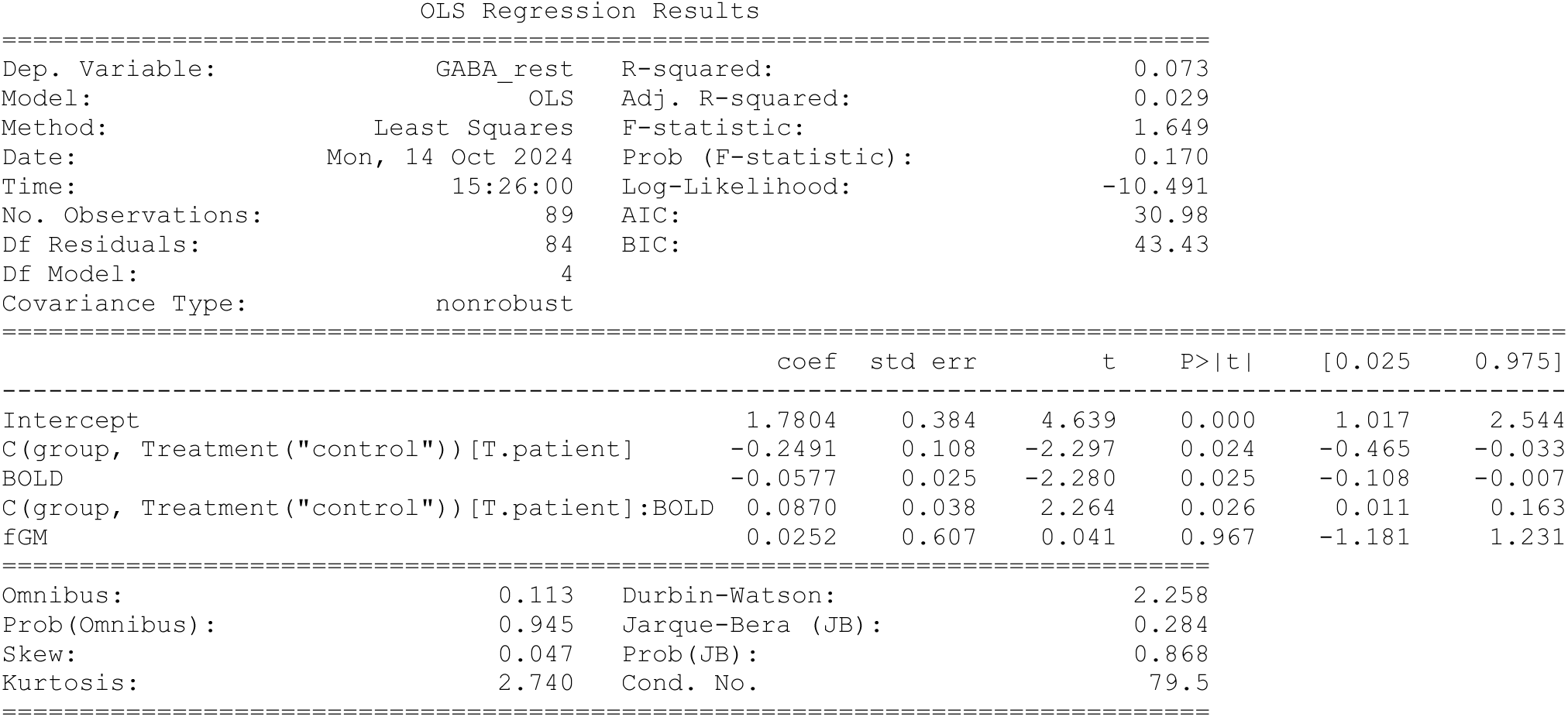

GABA+ estimate in relation to group (patient, control), task status (task-OFF, task-ON), with grey matter fraction as a covariate and subject as the grouping variable (ie, GABA ∼ C(group)*C(task_state) + fGM), after filtering observations with strong residuals:

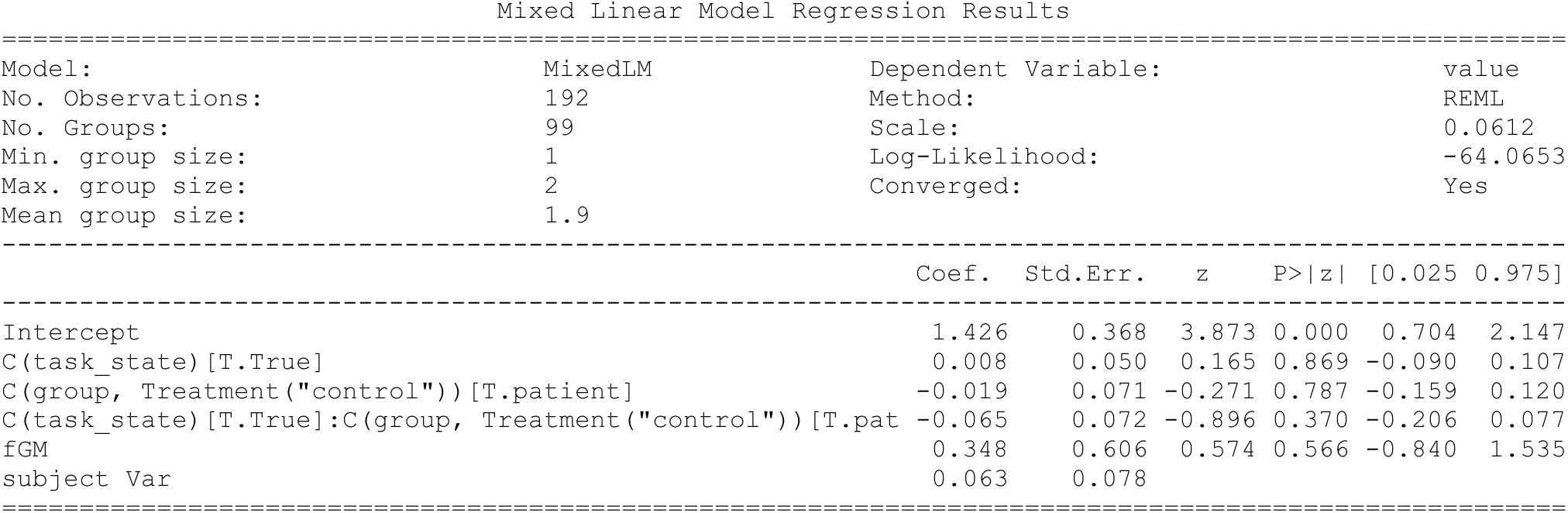

### C.3 Exploratory correlational tests

**Supplementary Table 5.**
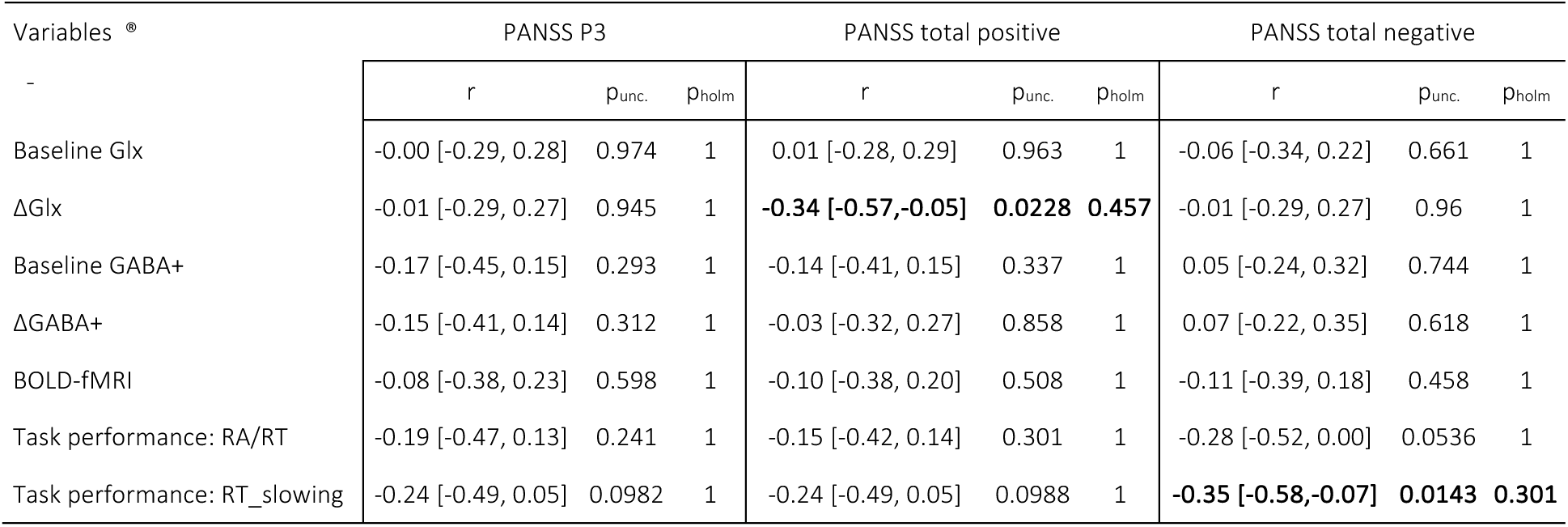
Exploratory correlational testing; skipped Spearman correlation with 95% confidence interval.

